# Transcriptional comparison of Testicular Adrenal Rest Tumors with fetal and adult tissues

**DOI:** 10.1101/2020.05.07.082313

**Authors:** Mariska A.M. Schröder, Fred C.G.J. Sweep, Antonius E. van Herwaarden, Alan E. Rowan, Darren Korbie, Rod T. Mitchell, Nike M.M.L. Stikkelbroeck, Hedi L. Claahsen – van der Grinten, Paul N. Span

**Affiliations:** Department of Pediatrics, Radboud Amalia Children’s Hospital, Radboud university medical center, 6500 HB Nijmegen, The Netherlands; Department of Laboratory Medicine, Radboud Institute for Molecular Life Sciences (RIMLS), Radboud university medical center, 6500 HB Nijmegen, The Netherlands; Australian Institute for Bioengineering and Nanotechnology, The University of Queensland, Brisbane Qld 4072, Australia; MRC Centre for Reproductive Health, University of Edinburgh, Edinburgh EH16 4TJ, United Kingdom; Department of Internal Medicine, Radboud University Medical Center, 6500 HB Nijmegen, The Netherlands; Radiotherapy & OncoImmunology Laboratory, Department of Radiation Oncology, Radboud Institute for Molecular Life Sciences (RIMLS), Radboud university medical center, 6500 HB Nijmegen, The Netherlands

**Keywords:** Adrenal, Congenital, Hyperplasia, TART, Tumor

## Abstract

Testicular Adrenal Rest Tumors (TART) are a common complication of unknown origin in patients with Congenital Adrenal Hyperplasia. These benign tumors may derive from cells of adrenal origin or from pluripotent progenitor cells from the fetal adrenogonadal primordium. By comparing the transcriptome of TART with fetal- and adult-testis and adrenal tissues, this study aims to unravel the origin of TART. Targeted transcriptome sequencing was followed by unsupervised clustering-, differential expression-, functional enrichment- and pathway analyses. Immunohistochemistry demonstrated co-expression of adrenal-specific CYP11B1 and testis-specific HSD17B3 in TART, indicating the existence of a distinct TART cell exhibiting both adrenal- and testicular characteristics. Principal component analysis indicated that the TART transcriptome was distinct from either adrenal or testis fetal tissue, making a progenitor-like phenotype of TART unlikely. Rather, TART appears to originate from -or differentiate into-a mature cell type, with both adrenal- and testicular characteristics. The present study, the first to describe the TART transcriptome, expands knowledge about the identity and functional characteristics of TART and identifies clinically targetable pathways associated with fibrosis.

## Introduction

Congenital Adrenal Hyperplasia (CAH) caused by defects in steroid 21-hydroxylase (CYP21A2) is an autosomal recessive disorder affecting adrenal steroid biosynthesis, leading to impaired glucocorticoid production (1). Consequently, due to a lack of negative feedback by glucocorticoids, adrenocorticotropin hormone (ACTH) production by the pituitary gland is increased, resulting in hyperplasia of the adrenal cortex (2). Steroid precursors upstream of the enzymatic defect accumulate and shunt into the non-affected adrenal androgen pathway resulting in elevated adrenal androgen production (3). Testicular tumors are a common complication in adult male CAH patients, with a prevalence ranging from 14% to 89% depending on the method of detection, patient age, and severity of the disease (4). These tumors are benign lesions, but because of their location within the rete testis and assumed mechanical obstruction of the seminiferous tubules, they may cause testicular damage and infertility. The tumors have histological and functional features of adrenal tissue (5) and, therefore, are called ‘testicular adrenal rest tumors’ (TART). However, the origin and etiological features of TART are still unclear.

Initially, TART was considered to originate from ectopic embryologic adrenal rests that descend along with the gonads during embryological development. It was thought that in CAH patients, elevated ACTH levels present *in utero* may prevent the regression of this ectopic adrenal tissue. Clark et al. (6) reported that TART tissue is homogeneous with morphological and biochemical characteristics of steroid-secreting adrenocortical cells with little similarity to normal testicular tissue. However, in addition to the expression of genes for adrenal-specific enzymes (*CYP11B1* and *CYP11B2*) and receptors (ACTH receptor *(MC2R)*; Angiotensin II receptor), testicular-specific gene expression patterns (*HSD17B3*; *LHCGR*; *GATA4*; possibly *INSL3*) have also been reported for TART (4). The presence of both adrenal- and testis-specific markers might indicate that TART arises from a common undifferentiated pluripotent progenitor, as the testis and adrenal gland are derived from the same adrenogonadal primordium. One of the candidate cell types are fetal Leydig cells (FLC), which have both adrenal- and testicular characteristics and are reported to be ACTH-sensitive (7). Alternatively, Val et al. (8) described the presence of an alternative adrenal-like cell type in the normal-developing mouse testis, which presents both adrenal- and Leydig-cell properties (hCG responsiveness) and suggests that testicular adrenal rests may develop from this cell population. TART appears to be already present in neonates (9) and infants (10), confirming the hypothesis of an embryological origin of TART.

ACTH is considered to be a growth promoting factor of TART, as the growth of TART is associated with poor hormonal control (11) and correlates with increased ACTH levels (12, 13). Besides, intensifying glucocorticoid treatment is sometimes effective in reducing tumor size, most likely due to the suppressing effects on ACTH levels (12, 14-16). However, an association between (long-term) hormonal control and the development or size of TART is not observed in all association studies (17, 18) and intensified glucocorticoid treatment does not result in tumor shrinkage in all cases (4). Longstanding tumors may become fibrotic (5), potentially explaining the tumors’ loss of responsiveness to glucocorticoid treatment (19) and the lack of correlation between long-term disease control and TART size in some studies. A better understanding of those fibrotic processes is required to prevent glucocorticoid-unresponsiveness and progression of TART.

Multiple studies have focused on the characterization of TART using an a priori selection of markers, but an unbiased characterization of TART is lacking. Knowledge of the identity and functional characteristics of TART will aid in the development of therapies to inhibit or prevent the development of TART. This study aims to unravel the identity and etiology of TART, by comparing the transcriptomic profiles of TART with both fetal- and adult-testis and adrenal tissues. By including both fetal and adult tissues in the (targeted) RNA sequencing panel, we aim to establish whether TART has a pluripotent progenitor phenotype or a more differentiated mature phenotype. We present the most robust characterization of TART to date, where the transcriptome of fourteen bilateral TART samples from seven CAH patients and two bilateral TART samples from a patient with recurrent Cushing’s disease is compared with the transcriptome of ‘normal’ adult adrenal-, adult testis-, fetal adrenal-, and fetal testis tissues.

## Results

### TART cells express adrenal- and Leydig cell-specific steroidogenic enzymes

Previous studies indicated the expression of adrenal- and testis-specific genes in TART (4). To study the spatial localization of adrenal- and testis-specific markers at the protein level, immunohistochemistry was performed. TART tissue sections were co-stained with Leydig cell-specific 17beta-Hydroxysteroid dehydrogenase 3 (HSD17B3) and adrenal-specific Cytochrome P450 11B1 (CYP11B1) (Figure 1). This allowed us to study whether TART is homogeneously composed of a single cell type expressing both testis- and adrenal-specific genes, or whether TART is composed of a mixture of different cell types. Normal adult adrenal tissue (HSD17B3-/CYP11B1+) and adult testis tissue (HSD17B3+/CYP11B1-) served as positive and as negative controls. The staining of TART samples clearly showed that most cells expressed adrenal-specific CYP11B1, whereas specific HSD17B3 staining was observed in a subset of CYP11B1+ TART cells. Interestingly, the HSD17B3 staining co-localized only with the CYP11B1 staining and no exclusive HSD17B3 staining (without CYP11B1) was observed in TART tissue. This suggests that the adrenal- and testicular features of TART do not represent a mixed composition of cells that express either adrenal- or testis-specific markers, but points to the existence of a distinct cell with both adrenal- and testis-like characteristics. TART is likely heterogeneously composed of different cell types (HSD17B3 positive and HSD17B3 negative), partly being this unique distinct CYP11B1+/HSD17B3+ cell type.

**Figure 1.**
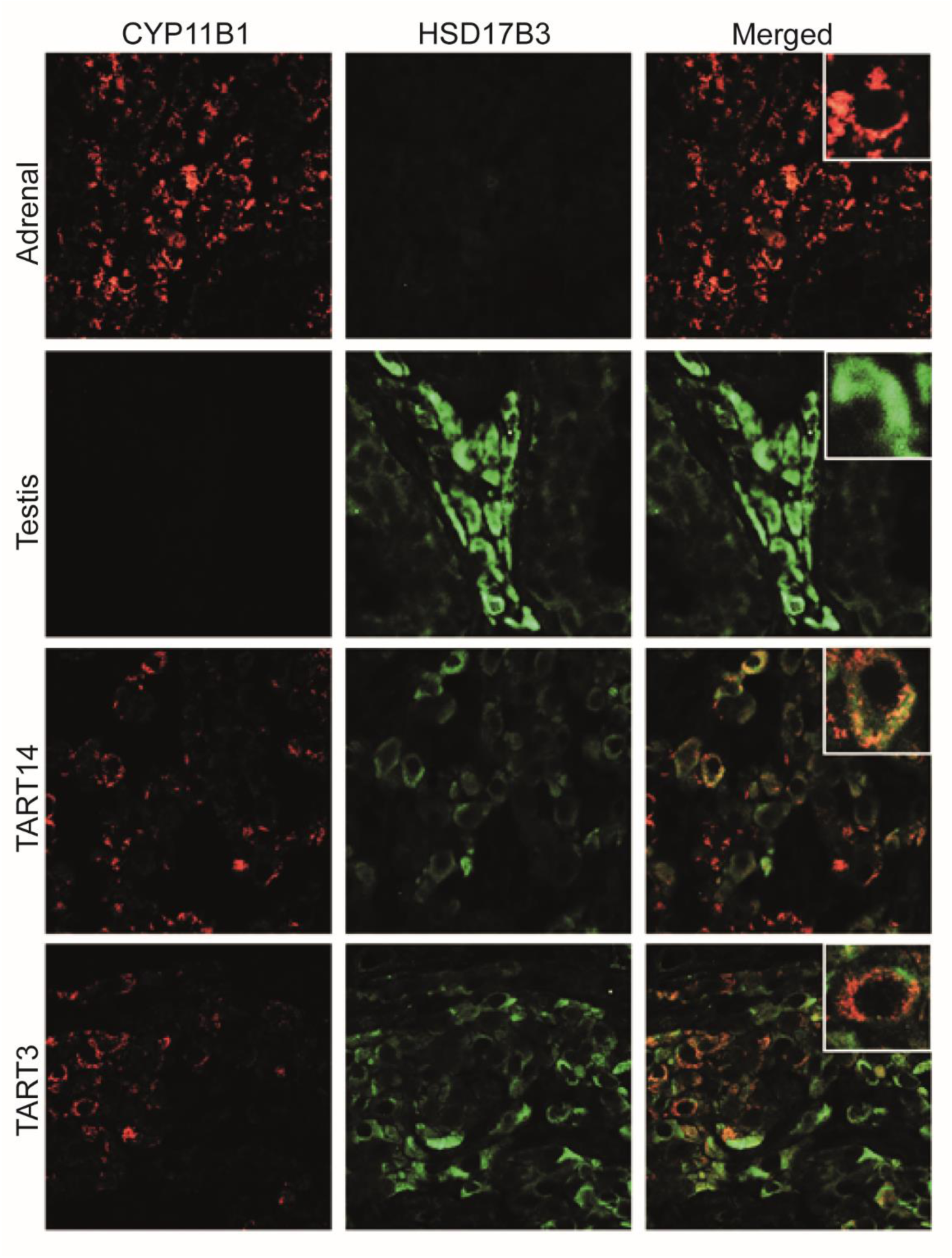
Double immunohistochemical staining of adrenal-specific 11β-hydroxylase (CYP11B1, red) and Leydig cell-specific 17β-hydroxysteroid dehydrogenase 3 (HSD17B3, green) on formaldehyde-fixed-paraffin-embedded (FFPE) material of adult adrenal, adult testis, and TART tissues.

### Transcriptome of TART is most similar to adult tissues

In order to study the (dis)similarity of TART with adult adrenal-, adult testis-, fetal adrenal-, and fetal testis tissues, targeted transcriptome sequencing was performed. Unsupervised hierarchical clustering using the complete clustering method showed that TART was most similar to either adult (adrenal- and testis) tissue, and least similar to either fetal (adrenal and testis) tissue (Figure 2A). Ten TART samples (TART1-5,8-12) from CAH patients and two TART samples that were removed from a patient with Cushing’s Disease (Cushing-TART (CT)) were clustered together with adult adrenal tissue, whereas four TART samples (TART6,7,13,14) were clustered together with adult testis tissue. Fetal adrenal and testis tissues were clustered separately. Principal component analysis (PCA) revealed that the different tissues were well clustered and effectively separated based on the top 500 high variance genes (Figure 2B). Minimal deviation was identified between bilateral TART samples obtained from the same patient. TART samples were least similar to fetal adrenal- and fetal testis tissues, whereas TART was most similar to adult adrenal samples, followed by adult testis tissues. K-means clustering of the PCA into five clusters (number of different tissues involved, assuming that the Cushing-TART (CT) samples are similar to the TART samples from CAH) showed two distinct clusters for fetal adrenal samples and fetal testis samples (Figure 2B). Most adult testis samples (9/10) were clustered separately. One adult testis sample was clustered together with six TART samples using k-means clustering but did not deviate from the other adult testis samples. The final cluster contained the other eight TART samples, together with the two Cushing-TART samples and all eleven adult adrenal samples. The four TART samples (TART6,7,13,14) that were clustered together with adult testis tissue by hierarchical clustering (figure 2A) showed least variance with regard to the testis samples compared to the other TART samples in the PCA plot (figure 2B). Thus, there was some variation between TART, where some TART samples showed less variance with regard to adult testis samples than other TART samples. In summary, based on unsupervised clustering methods, TART was most similar to adult adrenal tissue followed by adult testis tissue and least similar to fetal adrenal- and fetal testis tissue, making a pluripotent progenitor phenotype for TART unlikely.

**Figure 2.**
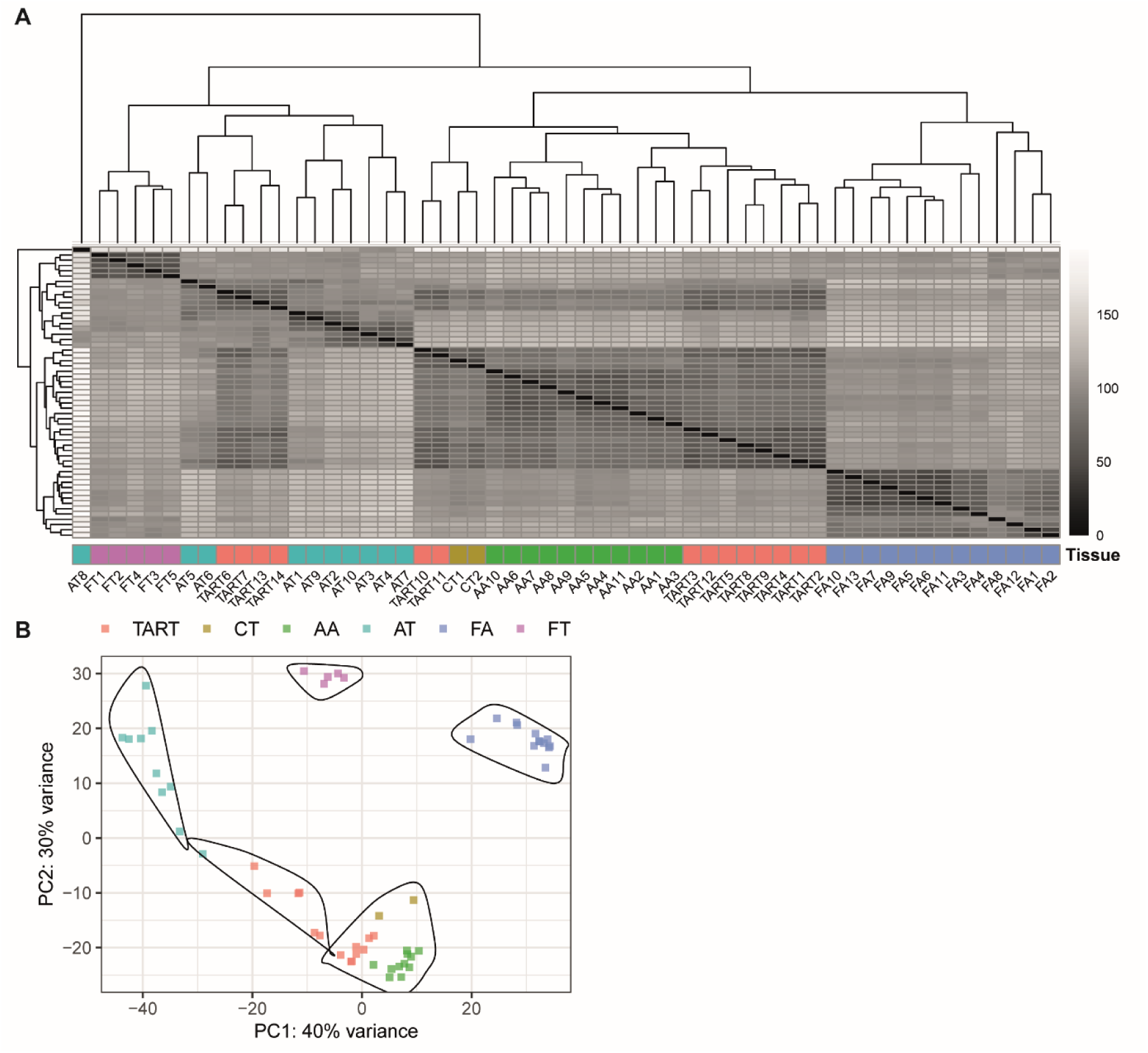
Hierarchical clustering analysis and Principal Component Analysis of TART and reference tissues. Abbreviations: CT = Cushing-TART; AA = Adult Adrenal; AT = Adult Testis; FA = Fetal Adrenal; FT = Fetal Testis. (A) Heatmap of unsupervised hierarchical clustering using the complete algorithm. (B) Principal Component Analysis (PCA) followed by k-means clustering (k=5) of Principal Component 1 (PC1) and Principal Component 2 (PC2).

### TART express both adrenal- and testis-specific genes

As TART seemed to be most similar to adult adrenal and testis tissues, TART-specific gene expression patterns and characteristics were studied in more detail by comparing the transcriptomic profile of TART with the transcriptomic profile of adult testis and adrenal tissues. Differential expression analysis using DESeq2 identified 1877 differentially expressed genes (DEGs) (fold-change > 2; adjusted p-value < 0.05) in TART compared to adult testis tissue, including 494 upregulated genes and 1383 downregulated genes (Figure 3A). Compared to adult adrenal tissue 728 genes were differentially expressed in TART, of which 449 genes were upregulated and 279 genes were downregulated in TART (Figure 3B).

**Figure 3.**
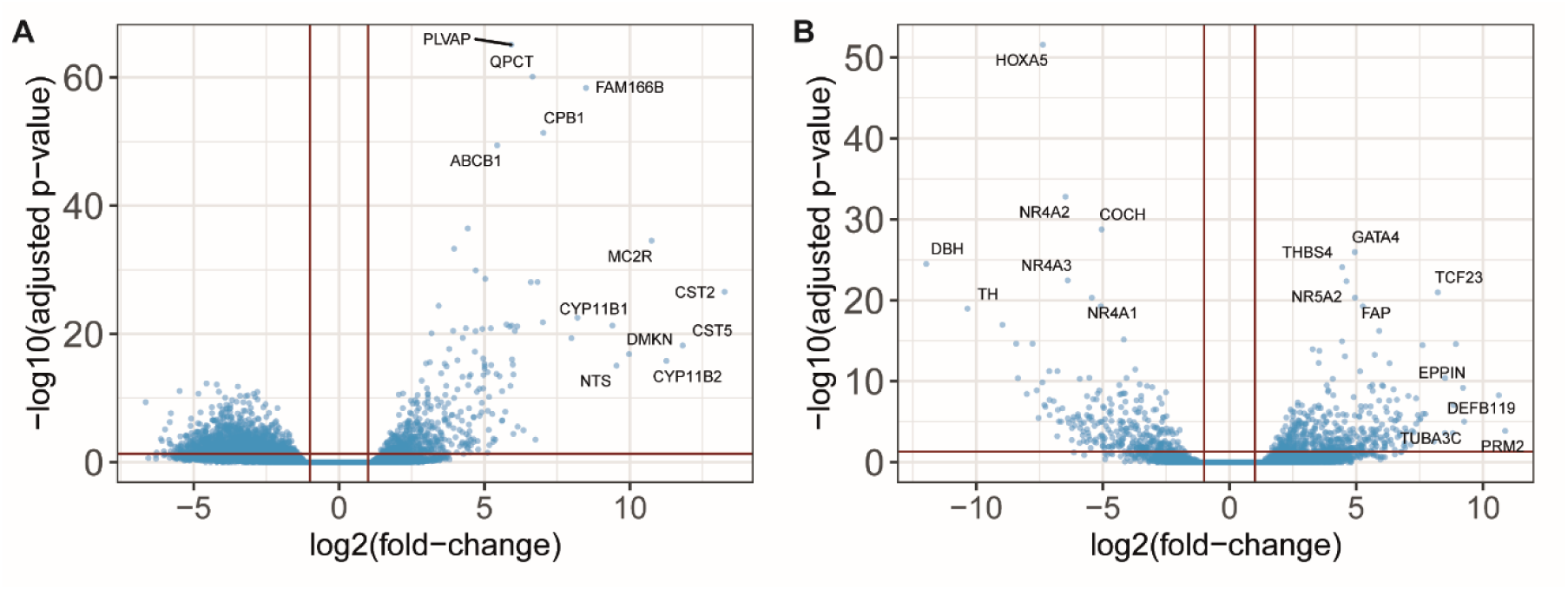
Volcano plots of differentially expressed genes in TART compared to adult testis tissue (A) and adult adrenal tissue (B). Vertical red lines indicate the log2 fold-change threshold at −1 and 1 and the horizontal line indicates the adjusted p-value threshold at 0.05.

Differential expression analysis showed high expression of the adrenal-specific genes *CYP11B1, CYP11B2*, and *MC2R* in TART versus testis tissue, suggesting adrenal specific features (Figure 3A), which is in line with previous research (4). Compared to adult adrenal tissue, TART highly overexpressed testis-specific *DEFB119, PRM2*, and *GATA4* (Figure 3B). *GATA4* expression in TART has been reported previously by our group (20). Previous studies proposed the expression of the testis-specific Luteinizing Hormone/Choriogonadotropin Receptor (*LHCGR*) in TART (16, 21). However, only very low expression levels were observed in this study and its expression did not deviate from that of the adrenal gland.

According to the Human Protein Atlas (HPA), 429 genes are expressed specifically in the testis (more than four-fold higher expressed compared to other analyzed tissues) (22, 23). At least 31 of those testis-specific genes were identified to be increased expressed in TART compared to adrenal tissue, suggesting testis-specific features of TART. Both Leydig cell-specific- (*INSL3*; *HSD17B3*) as well as germ-cell specific genes (e.g. *FATE1*; *DMRT1*; *DDX4*) (22, 24, 25) were overexpressed in TART compared to adult adrenal tissue. The overall expression of fourteen of those 31 testis-specific genes was higher in testis compared to TART tissue. It is worth noting that the expression of testis-specific genes in TART samples was variable (Supplementary Figure S1). The expression of testis-specific genes was high in the TART samples that deviated most from adrenal tissue with PCA (TART 6,7,12,13,14) but low in the other TART samples. The variability in testis-specific gene expression between TART samples was not associated with concentrations of ACTH, 17-hydroxyprogesterone, or androstenedione at the time of surgery, which have been presented previously (26). In summary, differential expression analysis highlighted the expression of both adrenal-specific genes, as well as testis-specific genes in TART. Whereas adrenal-specific genes were expressed in all TART samples, testis-specific gene expression patterns were variable in TART.

### Upregulated genes in TART are associated with steroidogenesis, fibrosis, and reproduction

Gene ontology (GO) overrepresentation analysis was performed to explore the potential function of the upregulated and downregulated DEGs in TART compared to adult testis and adrenal tissues. For the upregulated DEGs, GO analysis revealed a total of 375 and 112 significantly enriched GO terms for TART versus adult testis and adrenal tissues respectively. Subsequent filtering of redundant GO terms reduced the number to 172 and 42 enriched GO-terms for TART versus adult testis and adrenal tissues respectively (Supplementary Table 1 and 2). These GO term annotations are divided into three distinct categories; molecular functions (MF), biological processes (BP) and cellular compartments (CC). The five most significant enriched GO terms for each category are shown in Figure 4A and B.

**Figure 4.**
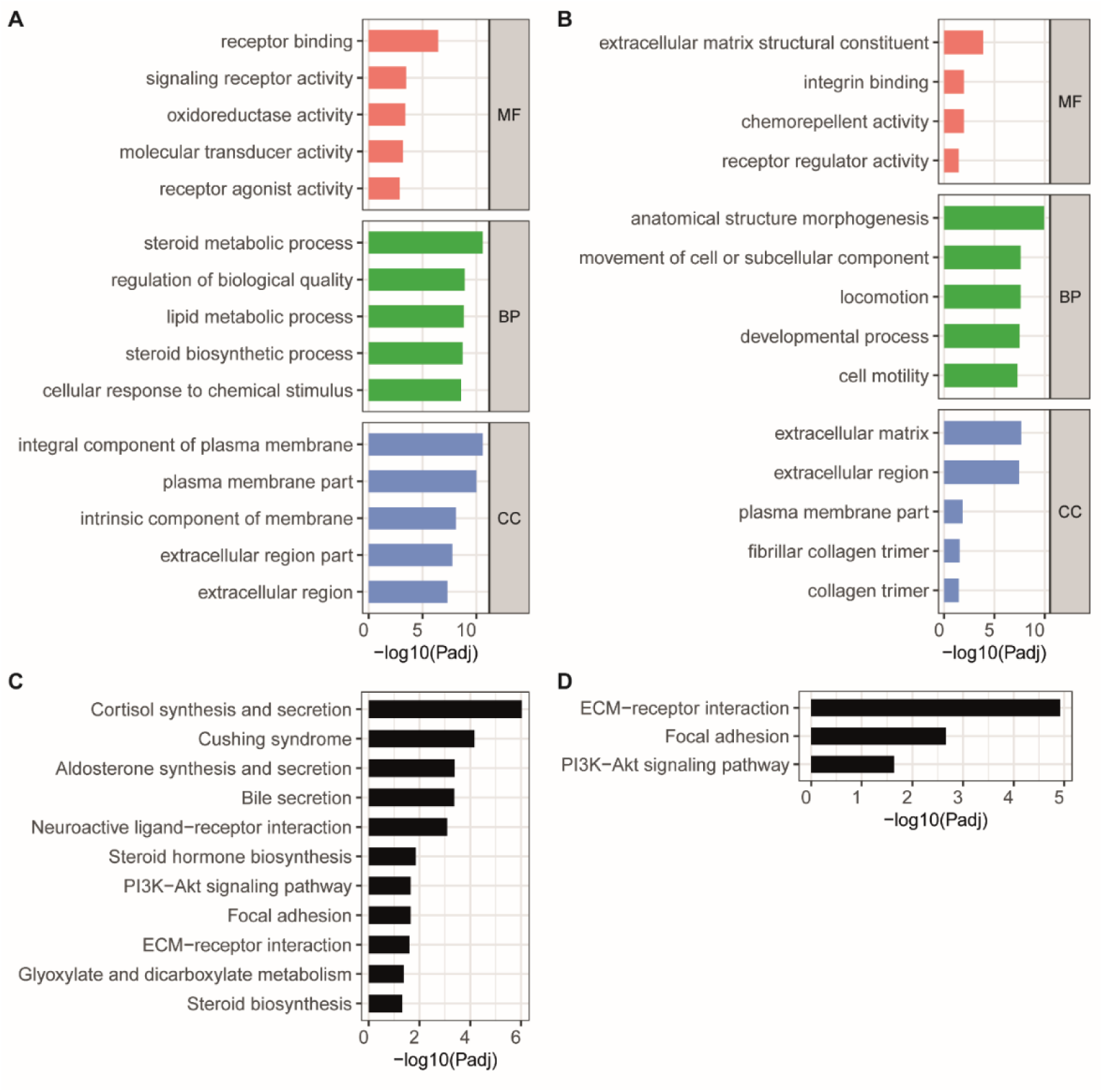
Functional overrepresentation and Kyoto Encyclopedia of Genes and Genomes (KEGG) pathway analysis were performed on significantly upregulated genes (fold-change > 2; adjusted p-value < 0.05) in TART compared to testis- (left panels) and to adrenal tissue (right panels). The top five most significant enriched Gene Ontology (GO) terms of each distinct category (molecular functions (MF), biological processes (BP) and cellular compartments (CC)) are shown for the overrepresented upregulated genes in TART versus adult testis (A) and TART versus adult adrenal tissue (B). All significant enriched KEGG pathways (Benjamini-Hochberg adjusted p-value < 0.05) are shown for TART versus adult testis (C) and TART versus adult adrenal tissue (D).

Overall, the upregulated genes in TART compared to adult testis tissue (Figure 4A) were associated with receptor binding/activity (MF) and the biological processes (BP) ‘steroid metabolic process’ and ‘cellular response to chemical stimulus’. The ‘plasma membrane’ and ‘extracellular region’ were, in general, the overrepresented cellular compartments (CC) (Supplementary Table 1). KEGG pathway analysis revealed overrepresentation of pathways associated with steroid hormone synthesis and secretion, as well as focal adhesion and ECM-receptor interaction (Figure 4C) in TART compared to adult testis tissue. Thus, both functional analysis and pathway analysis confirm steroid producing cell features of TART. The overrepresentation of the pathways ‘cortisol synthesis and secretion’ (Supplementary Figure S2) and ‘aldosterone synthesis and secretion’ (Supplementary Figure S3) highlighted the increased expression and activation of the ACTH receptor (MC2R) and angiotensin II receptor type I (AGTR1; AT1).

For the upregulated genes in TART compared to adult adrenal tissue (Figure 4B), the molecular functions ‘extracellular matrix structural constituent’ and ‘integrin binding’ were most highly enriched. The overexpression of germ-cell specific genes contributed to the overrepresentation of developmental- and reproductive processes (BP) (Supplementary Table 2). The CC category indicated overrepresentation of extracellular matrix (ECM)-related genes. KEGG pathway analysis revealed increased activity of the ECM-receptor interaction- and focal adhesion pathways in TART compared to adult adrenal tissue (Figure 4D). The overexpression of collagen genes and other extracellular matrix structural constituents (e.g. FBLN2; LAMA1) together with the increased expression of extracellular matrix metalloproteinases (e.g. ADAM12; ADAM23) and extracellular matrix remodeling genes (e.g. FAP) suggests active ECM remodeling in TART compared to adrenal tissue.

Next, TART-specific GO-terms were explored, as they may comprise actionable targets to prevent the development of TART. Thirty-nine genes were upregulated in TART compared to both adult testis and adrenal tissues. For those genes, three gene ontology terms were enriched, being: ‘extracellular region’, ‘extracellular space’, and ‘extracellular region part’. Closer examination of the upregulated genes associated with those GO terms suggested the overrepresentation of genes associated with ECM degradation/remodeling or fibrosis.

### Downregulated genes in TART are associated with reproduction and the nervous system

For the downregulated genes, gene ontology analysis revealed a total of 149 and 476 significantly enriched GO terms for TART versus adult testis (Figure 5A) and adrenal tissue (Figure 5B) respectively. Subsequent filtering of redundant GO terms reduced the number to 48 and 149 enriched GO-terms (Supplementary Table 3 and 4). ‘Microtubuli motor activity’ (MF), ‘reproductive process’ (BP), and ‘sperm part’ (CC) were the most highly enriched GO terms for each category among the genes downregulated in TART compared to adult testis tissue, likely pointing to spermatogenic properties of testis tissue compared to TART.

**Figure 5.**
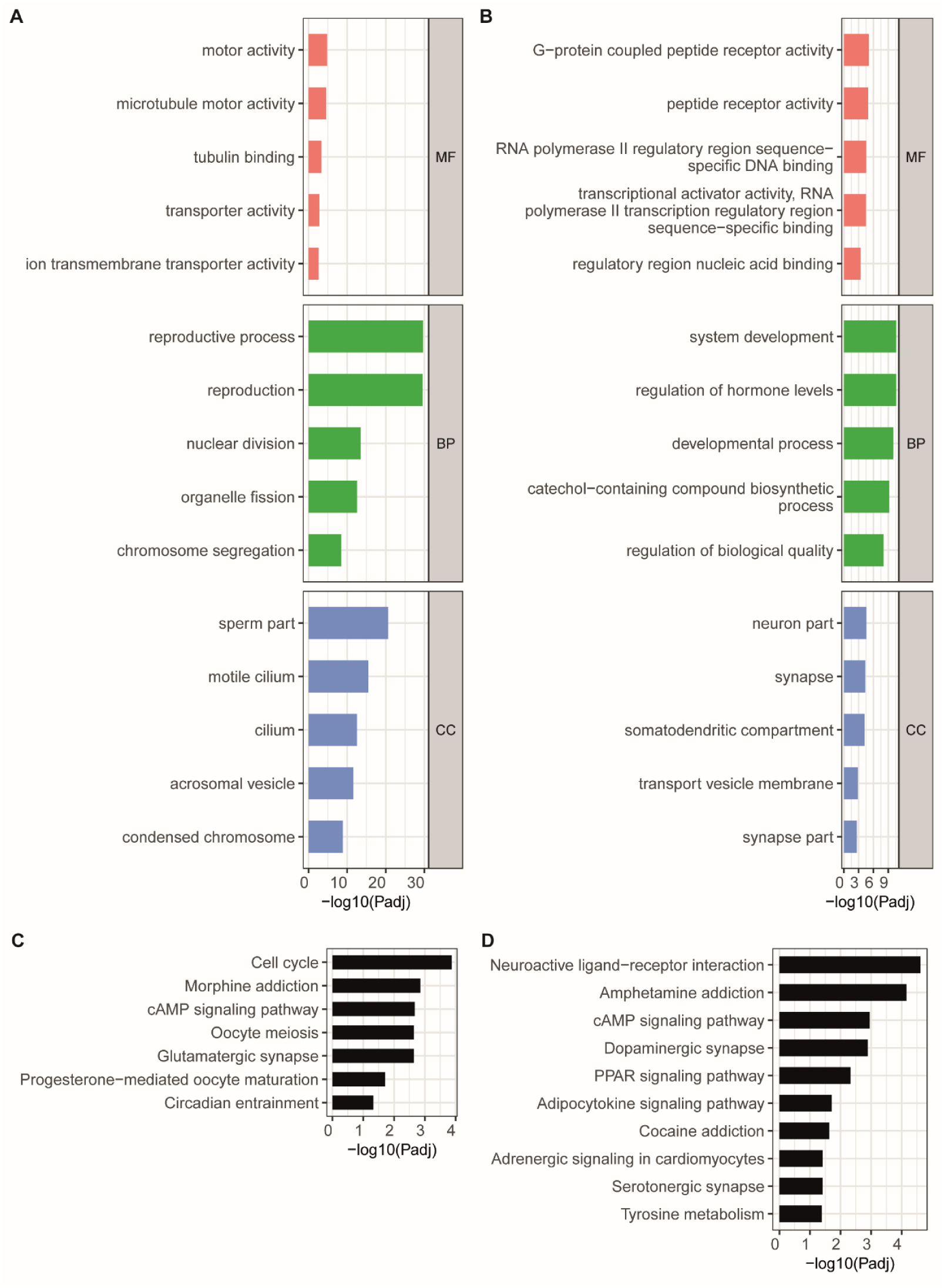
Functional overrepresentation and Kyoto Encyclopedia of Genes and Genomes (KEGG) pathway analysis were performed on significantly downregulated genes (fold-change < −2; adjusted p-value < 0.05) in TART compared to testis (left panels) and adrenal tissue (right panels). The top 5 most significant enriched Gene Ontology (GO) terms of each distinct category (MF, BP, CC) are shown for the overrepresented downregulated genes in TART versus adult testis (A) and TART versus adult adrenal tissue (B). All significant enriched KEGG pathways (Benjamini-Hochberg adjusted p-value < 0.05) are shown for TART versus adult testis (C) and TART versus adult adrenal tissue (D).

The genes downregulated in TART compared to adult adrenal tissue are associated with (among others) the molecular function ‘peptide receptor activity’. Overrepresented BPs are ‘system development’ and ‘regulation of hormone levels’, as well as GO-terms associated with the nervous system (Supplementary Table 4). Enriched cellular compartments comprised neuronal parts.

In summary, for the upregulated genes in TART compared to adult testis tissue, the functional annotations and pathways associated with steroid hormone biosynthesis were highly overrepresented. Gene ontology- and pathway analyses of the upregulated genes in TART compared to adult adrenal tissue, highlighted the overrepresentation of genes associated with the extracellular matrix and developmental processes/reproduction. Reproductive processes were, however, even higher represented by adult testis, compared to TART, as reported by the functional analysis of the downregulated genes in TART compared to testis tissue. Compared to adult adrenal tissue, TART showed lower expression of genes that are functionally associated with the nervous system.

## Discussion

TART is a common finding in adult male CAH patients that may lead to irreversible damage to the testis and infertility. The identity and functional characteristics of TART are incompletely understood. Previous research focusing on the characterization of TART reported both adrenal- (26-28) and testis-specific (20, 27) characteristics. However, these studies focused only on a subset of candidate markers. To our knowledge, this is the first study that characterizes TART using an unbiased approach. We present a unique and extensive transcriptome dataset of fourteen TART samples from seven CAH patients, two TART samples from a patient with recurrent Cushing’s Disease, eleven adult adrenal samples, ten adult testis samples, thirteen fetal adrenal samples, and five fetal testis samples. This study aimed to generate and use this dataset to study the identity and functional features of TART.

Because of previous identification of adrenal-and testicular markers in TART (4), it was speculated that TART might be derived from a more totipotent progenitor cell type, expressing both adrenal-specific markers as well as testis-specific markers. We proposed fetal Leydig cells as a potential candidate cell type, as they have both adrenal- and testicular characteristics and are reported to be ACTH-sensitive. Surprisingly, the TART transcriptome was least similar to fetal adrenal and testis tissues, making a ‘fetal’ progenitor-like phenotype of TART unlikely. The transcriptome of TART was most similar, but not equal to, adult adrenal samples, followed by adult testis samples, suggesting that TART cells either originate from or have differentiated into, a mature and unique distinctive cell type. TART cells do express both adrenal- and testis-specific markers. The co-expression of adrenal-specific CYP11B1 and Leydig cell-specific HSD17B3 within our samples suggests that the adrenal- and testicular features of TART do not represent a mixed composition of adrenal-like and testis-like cells, but points to the existence of a distinct TART cell with both adrenal- and testis-like characteristics. Nonetheless, Leydig cell-specific HSD17B3 was only expressed in a subset of TART cells, suggesting that TART might be heterogeneously composed of different cell types.

Our study emphasizes the high similarity between TART and adult *adrenal* tissue. The marked increased expression of the adrenal-specific genes *CYP11B1, CYP11B2* and *MC2R* in TART compared to testis tissue is in line with our previous research and underscores the robustness of our study (26). Functional analysis and KEGG pathway analysis highlight the adrenal steroidogenic properties of TART. Previous research indicated that TART produces adrenal-specific steroids (26, 29), as blood of CAH patients with TART present higher levels of adrenal specific 11-oxygenated androgens compared to blood of patients without TART (29). Furthermore, higher levels of adrenal-specific steroid hormones have been detected in spermatic vein sampling compared to peripheral blood samples in CAH patients (16, 26, 30). KEGG pathway analysis highlights the overexpression of genes involved in the adrenal steroid hormone biosynthesis and, more specifically, in cortisol and aldosterone synthesis. This suggests that TART has both zona fasciculata and zona glomerulosa-like characteristics. Cortisol biosynthesis is regulated by the hypothalamic-pituitary-adrenal axis and aldosterone secretion is regulated by the renin-angiotensin system. Differential expression and pathway analysis indicate the increased expression of the ACTH receptor (*MC2R*) and the Angiotensin II receptor (*AGTR1; AT1*) in TART compared to testis tissue. ACTH and angiotensin II have trophic effects on adrenocortical tissue (31, 32) and likely play a major role in the growth of TART (6, 27).

Our study also clearly demonstrates *testicular* characteristics of TART samples in addition to the adrenal characteristics. With PCA, TART samples were positioned between the adrenal tissue samples and testis tissue samples and next to adrenal-specific gene expression patterns. Testis-specific (i.e. genes reported to be specifically expressed in testis according to literature and the Human Protein Atlas) gene expression was also found in TART samples. Previous research proposed that the LHCGR receptor might play a role in the etiology of TART (16, 21). However, our data showed only very low numbers of normalized read counts for the LHCGR receptor in TART samples and no significant overexpression of the LHCGR receptor in TART compared to adult adrenal tissue was found, as the adrenal gland showed similarly low levels of *LHCGR* expression. This is consistent with previous findings, where we reported nearly equal expression levels of *LHCGR* in TART and adrenal tissue (27). Adrenocortical expression of low levels of *LHCGR* has been reported previously (33). Although expression levels of *LHCGR* were low, we did demonstrate expression of other testis-specific genes in TART. Remarkably, in addition to upregulation of genes specific for somatic testicular cells, we also identified upregulation of genes specifically expressed in spermatogonia, spermatocytes or spermatids. Subsequently, GO analysis demonstrated overexpression of genes associated with reproduction in TART compared to adult adrenal tissue. The expression of germ-cell-specific genes has not been reported in TART before and was unexpected, as TART has previously histologically been described as solely steroid-producing cells. The expression of testicular germ cell-specific genes in ectopic tissues has been widely described for cancer tissues (34). Because of the expression of both germ cell-specific genes and Leydig cell-specific genes, it may be argued that the expression of those testis-specific genes in TART is caused by the presence of surrounding testis tissue in the TART biopsies or that TART cells are intermingled with healthy Leydig cells and germ cells. However, the TART tissues removed during surgery were well-demarcated and ‘contamination’ of Leydig cells with exclusive HSD17B3 expression (no CYP11B1 expression) was not observed in TART using IHC (not at the borders of the tissue sections, nor within the tissue sections). The co-expression of adrenal-specific CYP11B1 and testis-specific HSD17B3 in TART tissue indicates the presence of a distinct and unique cell type with both adrenal- and testis-specific features. It should be mentioned that, whereas CYP11B1 was expressed in nearly all cells, HSD17B3 was only expressed in a subset of cells and at lower levels than in Leydig cells. In addition, we observed some variation between individual TART samples, as some TART samples were more similar to testis tissue and presented higher expression of testis-specific genes compared to other TART samples. The variation between TART samples, and more specifically between the TART samples of different patients, might explain the contradicting results of previous case reports where TART with (4) or without (6) testis-specific characteristics have been reported. Benvenga et al. (21) hypothesized that pluripotent ACTH-sensitive cells in the testis may differentiate into either an adrenal cell type or testicular cell type depending on their stimulus. The variation of testis-specific expression between TART samples was not associated with blood ACTH concentrations, measured at the time of surgery. This, however, was a single measurement of blood ACTH, and valuable information on long-term cumulative exposure of TART to ACTH is lacking. Thus, we report that the testis-specific features of TART are not likely to be caused by testis-contamination, but points to the presence of a distinct TART cell with both adrenal- and testicular features. Additional testis-specific markers, including germ-cell specific markers, should be investigated in future research to determine what portion of TART has both adrenal- and testicular features.

Although the shift of TART samples in the PCA plot towards the adult testis samples was clearly accompanied by increased expression of testis-specific markers in those particular samples, additional factors causing the variation between individual TART samples and adult adrenal samples could be considered. Firstly, adrenal medullary RNA was present in the sequenced adrenal samples, while TART samples do not contain adrenomedullary cells. The adrenal medulla contains transformed chromaffinergic cells and the presence of those cells in the adrenal tissue samples most likely explains the enrichment of ‘neuronal GO-terms’ in the group of downregulated genes in TART compared to adult adrenal tissue. Secondly, compared to adult adrenal tissue TART overexpressed genes associated with the extracellular matrix and functional enrichment analysis highlighted GO terms associated with ECM remodeling. However, compared to testis tissue, the increased activity of those processes in TART was less clear. In comparison to testis tissue, only the collagen gene *COL10A1* was significantly overexpressed. Nonetheless, the biological process ‘regulation of extracellular matrix disassembly’ and ‘glycosaminoglycan binding’ were overrepresented in TART compared to testis tissue. In addition, examination of genes that were upregulated in TART compared to adult adrenal tissue and testis tissue showed overrepresentation of genes associated with the ECM region. Therefore, the identified overexpressed genes and overrepresented processes might point towards the fibrotic processes in TART or breakdown and remodeling of surrounding ECM, allowing further growth of the tumor. The emergence of fibrosis may explain the unresponsiveness of TART to glucocorticoid treatment in CAH patients (5, 19). Further research with regard to those processes is required, as they may represent interesting drug targets to prevent the progression of TART. Unfortunately, despite the relatively large set of TART samples, the sample size was too small to perform gene correlation network analyses, which would allow for the identification of gene networks and their association with clinical parameters and pathological characteristics, such as fibrosis.

In summary, this study presents the most robust molecular characterization of TART to date and expands the knowledge about the identity and functional characteristics of TART. We report that TART is most similar to adult adrenal tissue, followed by adult testis tissue, and least similar to the fetal adrenal testis tissues, suggesting that TART originates from, or has differentiated into, a mature and distinct unique cell type, exhibiting both adrenal and testicular features. Future studies should focus on the factors causing intertumor/interpatient heterogeneity as well as on intratumor heterogeneity, using single-cell approaches.

## Materials and Methods

### Tissue collection

In total, sixteen TART samples were used for this study. Twelve bilateral TART samples from six CAH patients (mean age 27 year; range 23-32 year) were collected as previously described (26). Written informed consent was obtained from all patients. Another two frozen histologically proven TART samples from one anonymous CAH patient were obtained. Additionally, two bilateral TART samples were collected from a 27-year-old patient with recurrent Cushing’s Disease since the age of eleven (35). Those tumors have previously been characterized and showed similar characteristics to TART from CAH patients and were analyzed separately from the TART samples obtained from CAH patients. Frozen anonymous samples of eleven normal adult adrenal glands and ten normal adult testes were collected from the Pathology and Urology departments and used in accordance with the Code of Conduct of the Federation of Medical Scientific Societies in the Netherlands (http://www.federa.org/codes-conduct). The study was approved by the institutional review board: CMO Radboudumc #2016-2977 and CMO-nr 2004/007). In addition, five fetal testis samples and thirteen fetal adrenal samples (first and second trimester) were anonymously obtained following elective termination of pregnancy from the MRC Centre for Reproductive Health of the University of Edinburgh. Informed consent was obtained with ethical approval from the South East Scotland Research Ethics Committee (LREC08/S1101/1). The study is conformed to the principles set out in the WMA Declaration of Helsinki.

### Immunohistochemistry

Double immunofluorescence staining was performed on formalin-fixed paraffin-embedded material of TART, adult testis and adult adrenal tissue, obtained from the pathology department (Radboudumc). Five µm thick sections were deparaffinized in neoclear (sigma) and hydrated. Antigen retrieval was performed by heating the samples in Tris-EDTA buffer (pH 9) at 95°C for 30 minutes. Samples were blocked for 30 minutes with 5% normal donkey serum and incubated overnight with adrenal-specific CYP11B1 (1:200) and Leydig cell-specific HSD17B3 (13415-1-AP; Proteintech; 1:100). The monoclonal CYP11B1 antibody (hCYP11B1-80-2-2) was kindly provided by Dr. Gomez Sanchez (36). Samples were incubated for 90 minutes with Alexa Fluor 488-conjugated donkey-anti-rabbit secondary antibody (1:300; Invitrogen; A21206) and Cyanine Cy™3-conjugated donkey-anti-rat secondary antibody (1:300; Jackson ImmunoResearch; 712-166-150) at room temperature. Tissue sections were imaged at 40x on a Zeiss LSM900 using the Zeiss Zen 3.0 software. Brightness and contrast were equally adjusted in all images.

### RNA isolation, library preparation, and sequencing

RNA was isolated using the total RNA Purification Kit (Norgen Biotek) according to the manufacturer’s protocol. RNA quantity and purity were assessed using a NanoDrop 2000 Spectrophotometer. RNA was reversed transcribed using Superscript VILO cDNA synthesis kit (Thermo Fisher Scientific) using a C1000 Thermal cycler (Biorad), according to the manufacturer’s protocol. Libraries were constructed using the Ion AmpliSeq Transcriptome Human Gene Expression kit (Thermofisher). Templating and sequencing were performed with the Ion PI HiQ OT200 kit, and Ion PI HiQ sequencing kit using Ion PI V3 chips (Life Technologies) and a Proton Sequencer (Thermofisher). All RNA samples gave libraries of the expected size, with no observable amplification of no-template controls. Mapping of read data was performing automatically on the Torrent Suite server, against a reference fasta file composed of the transcript sequences of the target genes, and total read counts per target gene were then tallied.

### Unsupervised clustering analysis

Read counts were normalized using the R/Bioconductor package DESeq2 (R version 3.6.2) (37), which normalizes for both sequencing depth and RNA composition. The absence of batch effects was confirmed using Principal Component Analysis (PCA). Unsupervised hierarchical clustering (clustering method ‘complete’) was performed, after logarithmic transformation using the Variance Stabilizing Transformation (vst) function (DESeq2). PCA was performed on logarithmically transformed data of the top 500 genes with the highest variance over the different samples.

### Differential expression analysis and functional enrichment analysis

Differential expression analysis on the transcriptome data was performed using DESeq2. The threshold for differentially expressed genes was set at an absolute fold-change > 2. Functional enrichment analysis was performed using the R package GOseq (version 1.38.0) (38). GOseq compensates for the transcription length selection bias and is based on Wallenius non-central hypergeometric distribution. Another reason to select GOseq as the enrichment tool is that GOseq, like DAVID (39) and the R package gprofiler2 (40), allows the use of custom background. Upregulated and downregulated genes were separately tested against the background genes that were included in the RNA-sequencing panel. Redundant GO terms were identified based on pairwise semantic similarity and removed using REVIGO (41) at a cut-off of 0.7. KEGG pathway enrichment analysis of upregulated- and downregulated genes against background genes was performed using the R package clusterProfiler (version 3.14.3) (42). For differential expression analysis, functional enrichment analysis and KEGG pathway analysis, p-values were adjusted by the Benjamini-Hochberg method to correct for multiple testing and control for false-discovery rate (FDR). Adjusted p-values < 0.05 were considered statistically significant.

## Acknowledgements

We would like to thank Dr. M. Engels for her practical and logistic contributions to the transcriptome sequencing experiment. We would like to thank Prof. Dr. C.E. Gomez-Sanchez for providing the CYP11B1 antibody and H. Peters (Department of Radiation Oncology, Radboudumc) for his assistance with the immunohistochemical stainings. We acknowledge the Microscopy Imaging Centre (MIC) of the Radboudumc for the use of their facility. This work was partially supported by the Medical Research Council (MRC) grant (G1100357).

## Authors Contributions

F.S., H.C., P.S., and A.H. designed the study and directed the project. R.M and N.S. provided human tissues. A.R. and D.K. conducted the (targeted) RNA sequencing and processed the data. M.S. performed the immunohistochemical staining and the RNA sequencing data analysis. Interpretation of the data was done by M.S., F.S., H.C., P.S., and A.H.. M.S. wrote the manuscript and designed the figures with support of P.S., H.C., F.S. and A.H. All authors critically revised the manuscript.

## Conflict of interest

The authors declare that there is no conflict of interest.

## Data Availability

The RNA-sequencing dataset will be made freely available in a public database.

## Supplementary Figures

**Supplementary Figure S1:**
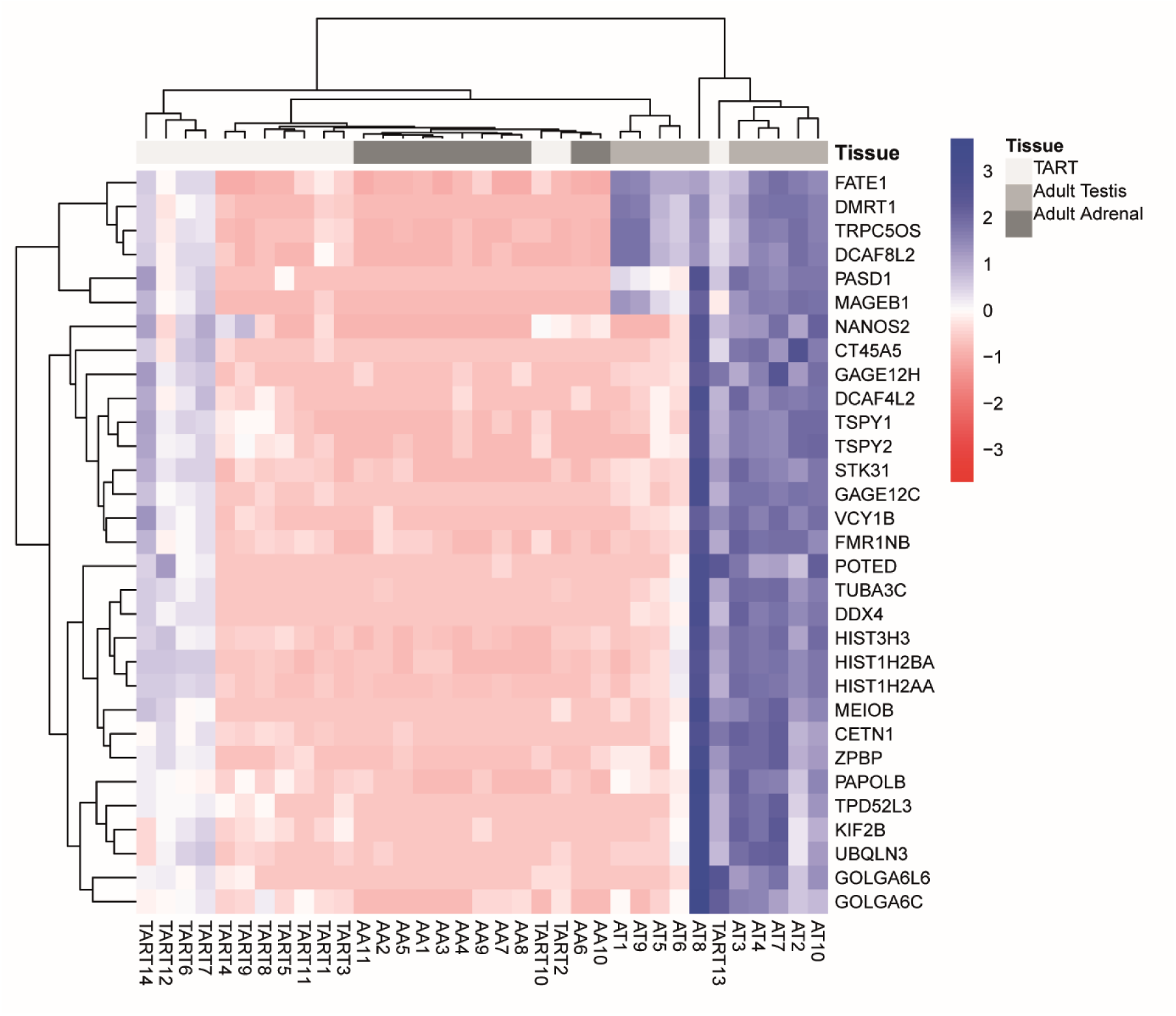
Heatmap showing relative log-transformed expression levels of 31 testis-specific genes that are significantly upregulated in TART compared to adult adrenal tissue. Blue color indicates relatively higher expression, red color indicates relatively lower expression compared to average expression.

**Supplementary Figure S2:**
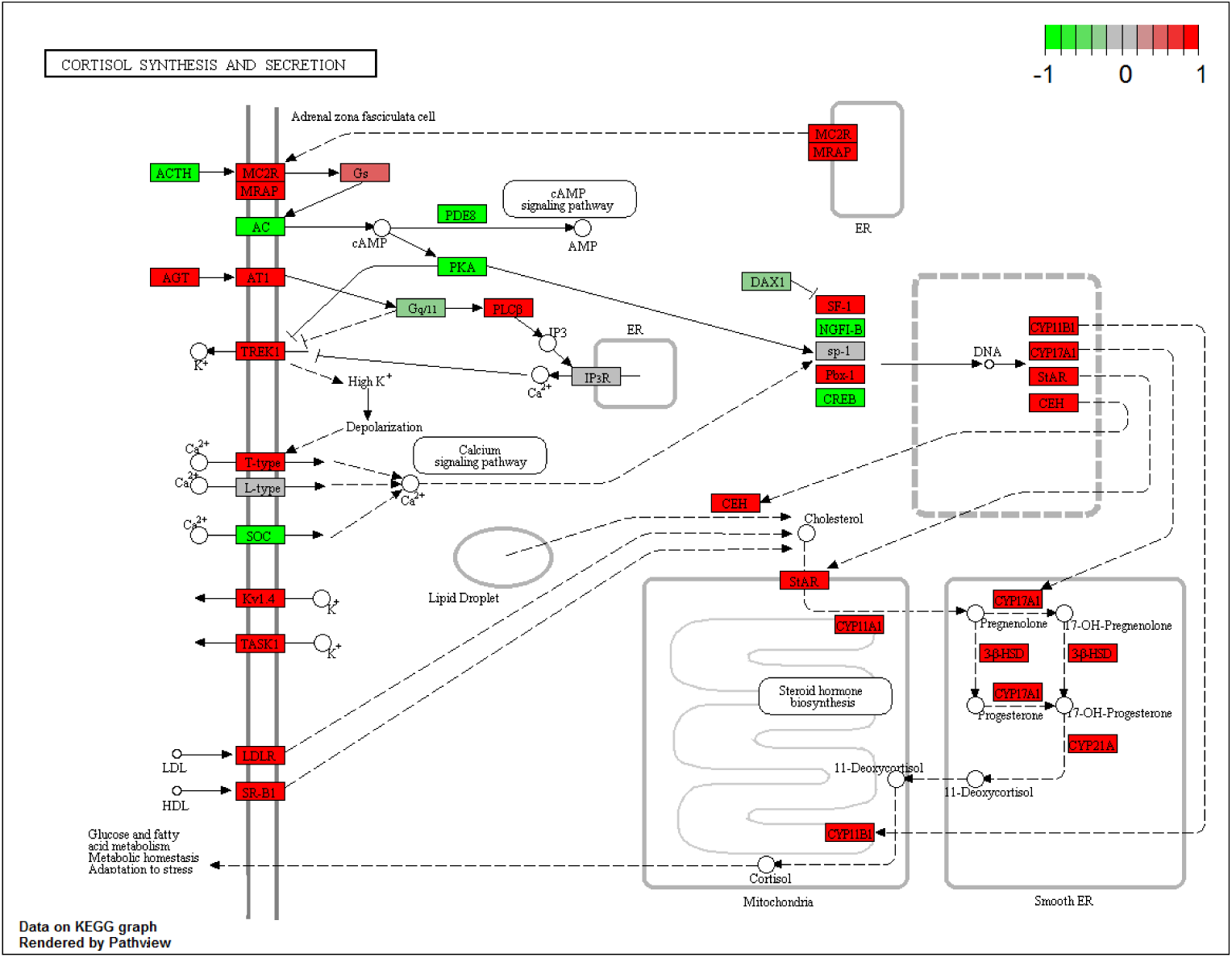
Pathway analysis (KEGG pathway) of expression levels in TART compared to adult testis tissue using the R package Pathview reveals overrepresentation of the cortisol synthesis and secretion pathway. Green color indicates the downregulated genes (log2 fold-change) and red color indicates upregulated genes.

**Supplementary Figure S3:**
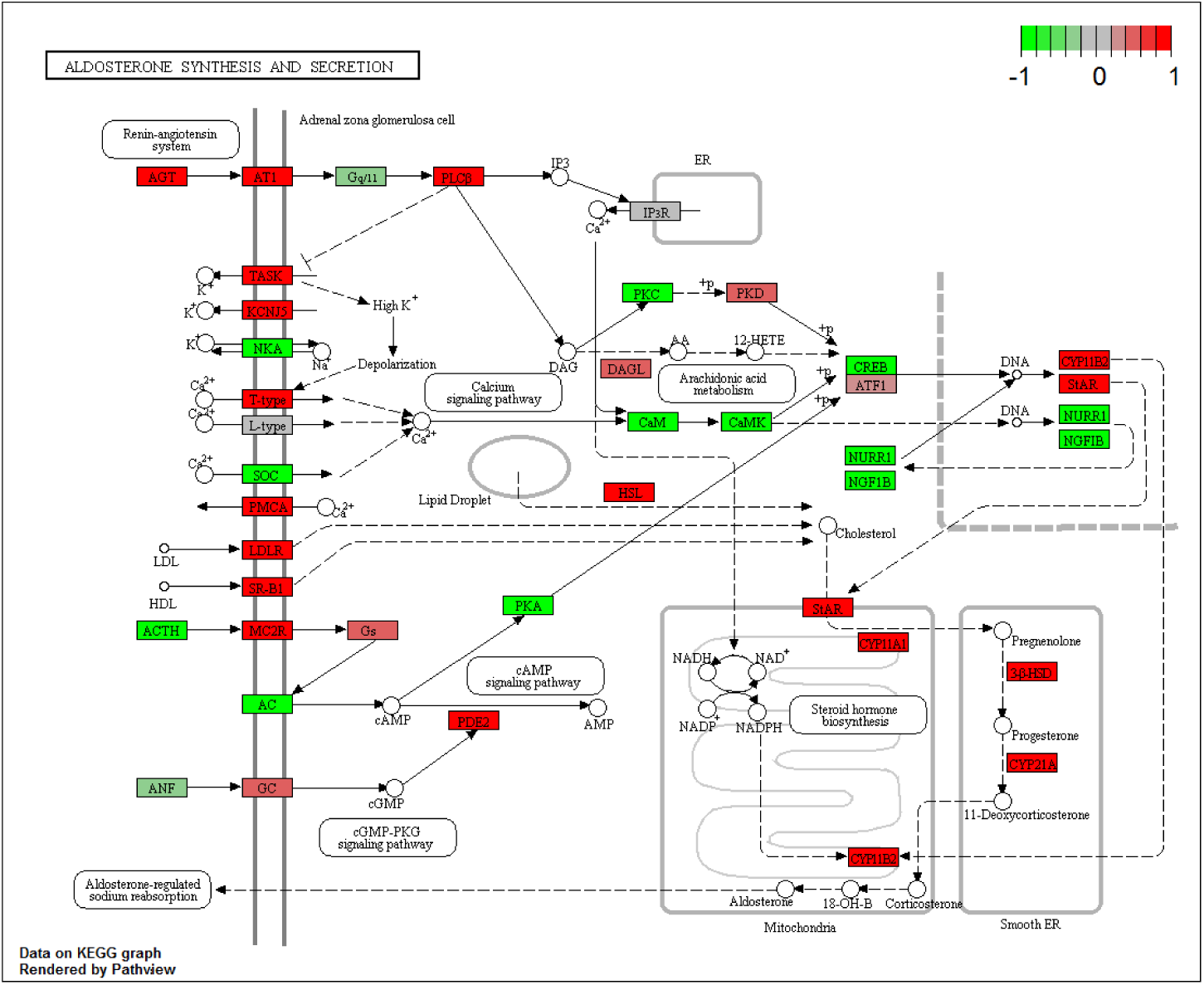
Pathway analysis (KEGG pathway) of expression levels in TART compared to adult testis tissue using the R package Pathview reveals the overrepresentation of the aldosterone synthesis and secretion pathway. Green color indicates the downregulated genes (log2 fold-change) and red color indicates upregulated genes.

## Notes

### Competing Interest Statement

The authors have declared no competing interest.

## References

1. El-Maouche D, Arlt W, Merke DP. Congenital adrenal hyperplasia. Lancet. 2017;390(10108):2194–210.

2. White PC, Speiser PW. Congenital adrenal hyperplasia due to 21-hydroxylase deficiency. Endocr Rev. 2000;21(3):245–91.

3. Speiser PW, White PC. Congenital adrenal hyperplasia. N Engl J Med. 2003;349(8):776–88.

4. Engels M, Span PN, van Herwaarden AE, Sweep F, Stikkelbroeck N, Claahsen-van der Grinten HL. Testicular Adrenal Rest Tumors: Current Insights on Prevalence, Characteristics, Origin, and Treatment. Endocr Rev. 2019;40(4):973–87.

5. Claahsen-van der Grinten HL, Otten BJ, Hermus AR, Sweep FC, Hulsbergen-van de Kaa CA. Testicular adrenal rest tumors in patients with congenital adrenal hyperplasia can cause severe testicular damage. Fertil Steril. 2008;89(3):597–601.

6. Clark RV, Albertson BD, Munabi A, Cassorla F, Aguilera G, Warren DW, et al. Steroidogenic enzyme activities, morphology, and receptor studies of a testicular adrenal rest in a patient with congenital adrenal hyperplasia. J Clin Endocrinol Metab. 1990;70(5):1408–13.

7. O’Shaughnessy PJ, Baker PJ, Johnston H. The foetal Leydig cell--differentiation, function and regulation. Int J Androl. 2006;29(1):90–5; discussion 105-8.

8. Val P, Jeays-Ward K, Swain A. Identification of a novel population of adrenal-like cells in the mammalian testis. Dev Biol. 2006;299(1):250–6.

9. Bouman A, Hulsbergen-van de Kaa C, Claahsen-van der Grinten HL. Prevalence of testicular adrenal rest tissue in neonates. Horm Res Paediatr. 2011;75(2):90–3.

10. Shanklin DR, Richardson AP, Jr., Rothstein G. Testicular Hilar Nodules in Adrenogenital Syndrome. The Nature of the Nodules. Am J Dis Child. 1963;106:243–50.

11. Mendes-Dos-Santos CT, Martins DL, Guerra-Junior G, Baptista MTM, de-Mello MP, de Oliveira LC, et al. Prevalence of Testicular Adrenal Rest Tumor and Factors Associated with Its Development in Congenital Adrenal Hyperplasia. Horm Res Paediatr. 2018;90(3):161–8.

12. Mazzilli R, Stigliano A, Delfino M, Olana S, Zamponi V, Iorio C, et al. The High Prevalence of Testicular Adrenal Rest Tumors in Adult Men With Congenital Adrenal Hyperplasia Is Correlated With ACTH Levels. Front Endocrinol (Lausanne). 2019;10:335.

13. Werneck G, Rodrigues EMR, Mantovani RM, Lane JSS, Silva IN. Testicular adrenal rest tumors in patients with congenital adrenal hyperplasia: 6 years of follow-up. J Pediatr Endocrinol Metab. 2019;32(5):519–26.

14. Armas-Alvarez AL, Salinas-Sanchez AS, Atienzar-Tobarra M, Virseda-Rodriguez JA. Testicular adrenal rest tumors. Rev Int Androl. 2018;16(3):128–30.

15. Claahsen-van der Grinten HL, Otten BJ, Sweep FC, Hermus AR. Repeated successful induction of fertility after replacing hydrocortisone with dexamethasone in a patient with congenital adrenal hyperplasia and testicular adrenal rest tumors. Fertil Steril. 2007;88(3):705 e5–8.

16. Combes-Moukhovsky ME, Kottler ML, Valensi P, Boudou P, Sibony M, Attali JR. Gonadal and adrenal catheterization during adrenal suppression and gonadal stimulation in a patient with bilateral testicular tumors and congenital adrenal hyperplasia. J Clin Endocrinol Metab. 1994;79(5):1390–4.

17. Reisch N, Scherr M, Flade L, Bidlingmaier M, Schwarz HP, Muller-Lisse U, et al. Total adrenal volume but not testicular adrenal rest tumor volume is associated with hormonal control in patients with 21-hydroxylase deficiency. J Clin Endocrinol Metab. 2010;95(5):2065–72.

18. Reisch N, Rottenkolber M, Greifenstein A, Krone N, Schmidt H, Reincke M, et al. Testicular adrenal rest tumors develop independently of long-term disease control: a longitudinal analysis of 50 adult men with congenital adrenal hyperplasia due to classic 21-hydroxylase deficiency. J Clin Endocrinol Metab. 2013;98(11):E1820–6.

19. Jha S, El-Maouche D, Marko J, Mallappa A, Veeraraghavan P, Merke DP. Individualizing Management of Infertility in Classic Congenital Adrenal Hyperplasia and Testicular Adrenal Rest Tumors. J Endocr Soc. 2019;3(12):2290–4.

20. Engels M, Span PN, Mitchell RT, Heuvel J, Marijnissen-van Zanten MA, van Herwaarden AE, et al. GATA transcription factors in testicular adrenal rest tumours. Endocr Connect. 2017;6(8):866–75.

21. Benvenga S, Smedile G, Lo Giudice F, Trimarchi F. Testicular adrenal rests: evidence for luteinizing hormone receptors and for distinct types of testicular nodules differing for their autonomization. Eur J Endocrinol. 1999;141(3):231–7.

22. Human Protein Atlas [Available from: http://www.proteinatlas.org.

23. Uhlen M, Fagerberg L, Hallstrom BM, Lindskog C, Oksvold P, Mardinoglu A, et al. Proteomics. Tissue-based map of the human proteome. Science. 2015;347(6220):1260419.

24. Djureinovic D, Fagerberg L, Hallstrom B, Danielsson A, Lindskog C, Uhlen M, et al. The human testis-specific proteome defined by transcriptomics and antibody-based profiling. Mol Hum Reprod. 2014;20(6):476–88.

25. Kim JY, Jung HJ, Yoon MJ. VASA (DDX4) is a Putative Marker for Spermatogonia, Spermatocytes and Round Spermatids in Stallions. Reprod Domest Anim. 2015;50(6):1032–8.

26. Claahsen-van der Grinten HL, Otten BJ, Sweep FC, Span PN, Ross HA, Meuleman EJ, et al. Testicular tumors in patients with congenital adrenal hyperplasia due to 21-hydroxylase deficiency show functional features of adrenocortical tissue. J Clin Endocrinol Metab. 2007;92(9):3674–80.

27. Smeets EE, Span PN, van Herwaarden AE, Wevers RA, Hermus AR, Sweep FC, et al. Molecular characterization of testicular adrenal rest tumors in congenital adrenal hyperplasia: lesions with both adrenocortical and Leydig cell features. J Clin Endocrinol Metab. 2015;100(3):E524–30.

28. Lottrup G, Nielsen JE, Skakkebaek NE, Juul A, Rajpert-De Meyts E. Abundance of DLK1, differential expression of CYP11B1, CYP21A2 and MC2R, and lack of INSL3 distinguish testicular adrenal rest tumours from Leydig cell tumours. Eur J Endocrinol. 2015;172(4):491–9.

29. Turcu AF, Mallappa A, Elman MS, Avila NA, Marko J, Rao H, et al. 11-Oxygenated Androgens Are Biomarkers of Adrenal Volume and Testicular Adrenal Rest Tumors in 21-Hydroxylase Deficiency. J Clin Endocrinol Metab. 2017;102(8):2701–10.

30. Bercovici JP, Fiet J, Gibault L, Volant A, Abalain JH, Floch HH, et al. Testicular adrenal rest tumours in salt wasting congenital adrenal hyperplasia (in vivo and in vitro studies). J Steroid Biochem Mol Biol. 2005;93(1):67–72.

31. Lotfi CF, de Mendonca PO. Comparative Effect of ACTH and Related Peptides on Proliferation and Growth of Rat Adrenal Gland. Front Endocrinol (Lausanne). 2016;7:39.

32. McEwan PE, Vinson GP, Kenyon CJ. Control of adrenal cell proliferation by AT1 receptors in response to angiotensin II and low-sodium diet. Am J Physiol. 1999;276(2):E303–9.

33. Bernichtein S, Alevizaki M, Huhtaniemi I. Is the adrenal cortex a target for gonadotropins? Trends Endocrinol Metab. 2008;19(7):231–8.

34. Fratta E, Coral S, Covre A, Parisi G, Colizzi F, Danielli R, et al. The biology of cancer testis antigens: putative function, regulation and therapeutic potential. Mol Oncol. 2011;5(2):164–82.

35. Puar T, Engels M, van Herwaarden AE, Sweep FC, Hulsbergen-van de Kaa C, Kamphuis-van Ulzen K, et al. Bilateral Testicular Tumors Resulting in Recurrent Cushing Disease After Bilateral Adrenalectomy. J Clin Endocrinol Metab. 2017;102(2):339–44.

36. Gomez-Sanchez CE, Qi X, Velarde-Miranda C, Plonczynski MW, Parker CR, Rainey W, et al. Development of monoclonal antibodies against human CYP11B1 and CYP11B2. Mol Cell Endocrinol. 2014;383(1-2):111–7.

37. Love MI, Huber W, Anders S. Moderated estimation of fold change and dispersion for RNA-seq data with DESeq2. Genome Biol. 2014;15(12):550.

38. Young MD, Wakefield MJ, Smyth GK, Oshlack A. Gene ontology analysis for RNA-seq: accounting for selection bias. Genome Biology. 2010;11(2).

39. Huang da W, Sherman BT, Lempicki RA. Systematic and integrative analysis of large gene lists using DAVID bioinformatics resources. Nat Protoc. 2009;4(1):44–57.

40. Raudvere U, Kolberg L, Kuzmin I, Arak T, Adler P, Peterson H, et al. g:Profiler: a web server for functional enrichment analysis and conversions of gene lists (2019 update). Nucleic Acids Res. 2019;47(W1):W191–W8.

41. Supek F, Bosnjak M, Skunca N, Smuc T. REVIGO summarizes and visualizes long lists of gene ontology terms. PLoS One. 2011;6(7):e21800.

42. Yu G, Wang LG, Han Y, He QY. clusterProfiler: an R package for comparing biological themes among gene clusters. OMICS. 2012;16(5):284–7.

